# Surface-based Characterization of Gastric Anatomy and Motility using Magnetic Resonance Imaging and Neural Ordinary Differential Equation

**DOI:** 10.1101/2022.10.17.512633

**Authors:** Xiaokai Wang, Jiayue Cao, Kuan Han, Minkyu Choi, Yushi She, Ulrich Scheven, Recep Avci, Peng Du, Leo K. Cheng, Madeleine R Di Natale, John B Furness, Zhongming Liu

## Abstract

Gastrointestinal magnetic resonance imaging (MRI) provides rich spatiotemporal data about the volume and movement of the food inside the stomach, but does not directly report on the muscular activity of the stomach itself. Here we describe a novel approach to characterize the motility of the stomach wall that drives the volumetric changes of the gastric content. In this approach, a surface template was used as a deformable model of the stomach wall. A neural ordinary differential equation (ODE) was optimized to model a diffeomorphic flow that ascribed the deformation of the stomach wall to a continuous biomedical process. Driven by this diffeomorphic flow, the surface template of the stomach progressively changes its shape over time or between conditions, while preserving its topology and manifoldness. We tested this approach with MRI data collected from 10 Sprague Dawley rats under a lightly anesthetized condition. Our proposed approach allowed us to characterize gastric anatomy and motility with a surface coordinate system common to every individual. Anatomical and motility features could be characterized for each individual, and then compared and summarized across individuals for group-level analysis. As a result, high-resolution functional maps were generated to reveal the spatial, temporal, and spectral characteristics of muscle activity as well as the coordination of motor events across different gastric regions. The relationship between muscle thickness and gastric motility was also evaluated throughout the entire stomach wall. Such a structure-function relationship was used to delineate two distinctive functional regions of the stomach. These results demonstrate the efficacy of using GI-MRI to measure and model gastric anatomy and function. This approach described herein is expected to enable non-invasive and accurate mapping of gastric motility throughout the entire stomach for both preclinical and clinical studies.

## I. INTRODUCTION

THE stomach is a hollow organ with thin layers of smooth muscle cells [1]. The musculated stomach wall forms a smooth and convoluted surface enclosing the intragastric ingesta. Its shape and movement exhibits gastric motor events that vary greatly across different gastric regions or different periods during digestion. The temporal dynamics and spatial coordination of gastric motor events are central to how the stomach stores, grinds, and eventually propels food at an efficient rate for digestion [2].

Many motility tests are available in the preclinical and clinical settings for studying and evaluating gastric motor functions in-vivo [3]–[5]. However, these tests often rely on invasive procedures to measure muscular activity confined to limited locations or time. The lack of sufficient spatial or temporal resolution or spatial coverage makes such measures inadequate for precise or high-throughput quantification for different aspects of gastric motility, such as gastric accommodation, phasic contractions, and coordination. Methods for in vitra motility assessment are valuable [6] but not easily applicable to in vivo settings or translable to humans.

Magnetic resonance imaging (MRI) has been increasingly used to image the stomach and characterize its anatomy and motility [7]–[11]. When an animal or human subject consumes a meal labeled with contrast agents, such as gadolinium [12] or manganese [13], [14], the luminal content of the stomach becomes visible with MRI and shows enhanced contrast against the surrounding tissues or organs. Changes in its size and shape are indicative of gastric emptying and motility, respectively. Using this strategy, recent studies demonstrated the unique potential of using MRI to study gastric physiology and pathophysiology and evaluate different treatments of gastric diseases in both preclinical and clinical settings [15]–[17].

Despite advances in gastric MRI, motility analysis has been primarily focused on the dynamic volumetric assessment of the food in the stomach or small bowel [12], [13], [18]–[21], as opposed to the 3-D movements of the wall. The volumetric analysis compresses 3-D shapes into 2-D, emphasizing the temporal evolution of the cross-sectional areas. The spatial variation along the circumferential orientation and their co-ordination are thus overlooked. In addition, results from the volumetric analysis are hard to generalize because the size and shape of the stomach change through an extensive range with the state of feeding and the nature of the diet across individuals. The volumetric analysis reports on the movement of the intragastric content, instead of the muscular activity of the stomach itself. It is thus not straightforward to relate the results of volumetric analysis to the underlying structural information pertaining to the stomach wall.

A more direct assessment of gastric motor events is to assess the biomechanical dynamics of the stomach wall that encloses and moves the luminal content. When the stomach distends itself to accommodate food, the stomach wall can be viewed as a 3-D surface, since its thickness is negligible relative to the much larger size of the stomach or the luminal volume. The surface of the stomach changes its shape dynamically due to a cascade of electrical, chemical, and mechanical processes, which are coupled to drive muscular activity and mechanical forces through the stomach wall. The luminal content acts passively to respond to the forces of shearing, squeezing, and grinding, appearing as the changes in the shape (surface) over time. These changes related to different gastric function could be better characterized by surface analysis, than the volumetric analysis. Therefore, tracking this temporal evolution of the spatial deformation of the surface is a more direct measurement of gastric motor events.

Furthermore, the 3-D surface-based analysis allows the extracted functional features and structural knowledge overlaid on the common template, facilitating the direct comparison between the two. In line with this idea, a generic surface template of the stomach (namely the scaffold) has been established to map data about gastric anatomy in rats [22]. Onto the scaffold, researchers have registered descriptions of the musculature [1], vasculature [23], enteric neurons [24], and vagal projections [25]. This scaffold is well suited to serve as a common representation for characterizing gastric anatomy and motility. If the scaffold can be shared and preserved between animals, conditions, and times, the structural and functional representations on the scaffold can be readily compared, summarized, and correlated at both individual and group levels.

For this surface-based analysis, the central task is to track the deformation of the surface enclosing the gastric volume. From the perspective of bio-mechanics, the deformation should preserve the integrity (connectivity) of the muscles and thus avoid intersections between any two areas of the surface. An algorithm that performs this task should also be subject to a constraint that all points on the surface should preserve their topology given any physiologically plausible deformation. Without preserving the topology and manifoldness, surface registration or deformation may over-estimate gastric motility or fail to be physiologically plausible, while a regularization can be used to favor, but not guarantee, a smooth and non-intersecting surface [14].

Here, we report a new and effective approach for characterizing gastric anatomy and motility using MRI and neural ordinary differential equation (ODE). In this approach, the neural ODE [26], [27] is used to model a diffeomorphic flow for morphing a topologically fixed surface template (i.e., scaffold) toward individualized and time-resolved surfaces that enclose the luminal volume observed with dynamic and contrast-enhanced MRI [12], [28]. The resulting surfaces describe the state-dependent anatomy and time-varying movement of the stomach with a surface coordinate system common for all animals and conditions. We developed this approach with MRI data from 10 healthy rats and tested its utility in uncovering the spatial, temporal, and spectral characteristics of gastric motor events and linking the resulting functional characteristics to the muscle thickness [1] co-registered on the same scaffold [29]. In the subsequent subsections, we first describe the intuition and theory that motivates our method, demonstrate its efficacy using simplified 2-D examples, then apply it to experimental data, characterize the gastric anatomy and motility in individual and group levels, and lastly demonstrate its potential in addressing the structure-function relationship by correlating the motility characteristics to the muscle thickness.

## II. METHODS

### A. Intuition and theory

When we model the stomach as a deformable surface, while ignoring the thickness of the stomach wall, the surface should be subject to its inherent structural, physical, and physiological constraints. That is, the surface of the stomach should be closed and smooth at any time. The movement or deformation of the stomach results directly from contraction or relaxation of muscles, which are topologically connective tissues that reside in the stomach wall. Their bio-mechanical responses are also continuous as mediated by the underlying physiology [1], [30], [31]. As such, the deformation must preserve its original topology, and be differentiable at any location, whereas it must not cause the surface to intersect at any time.

To satisfy these constraints, we model a diffeomorphic flow that drives every point on a surface to change its coordinate continuously. The diffeomorphic flow can be modeled by an ODE as generally expressed in

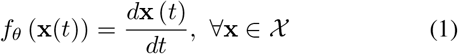

where x(*t*) is a vector representing the position of a moving point in space, *d*x(*t*)/*dt* expresses how this point moves from its position x at time t as a Lipschitz-continuous function of the position, *f_θ_* (x); its parameters are denoted as *θ*, and its domain is denoted as *χ*. This function governs how the surface as a whole changes its shape over time while every point on the surface flows along a continuous and unique path. If the diffeomorphic flow is described as a learnable and nonlinear function (e.g., a neural network), it can learn to explain complex movement or deformation given the positional data observed on the surface. This notion is utilized in a class of algorithms for computational anatomy [32], [33].

A diffeomorphic flow has important implications for our purpose. Equation (1) implies that *d*x(*t*)*/dt* always exists and is unique for any location x in the domain *χ* [34], [35]. Once a model of diffeomorphic flow is parameterized, it assigns a unique path to any point moving in the domain. It means that any two points never collide into each other as they move in the domain. If the diffeomorphic flow is applied to a closed and smooth surface, it morphs the surface while the surface always remains closed and smooth without any intersection. These properties represent the inductive bias for any model of diffeomorphic flow, regardless of its parameters.

In the context of our application, a model of diffeomorphic flow describes a continuous process of morphing a surface over a period of time. If the period starts from *t*_0_ and ends at *t*_1_, one can use an ODE solver to compute the morphing 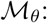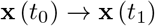 as

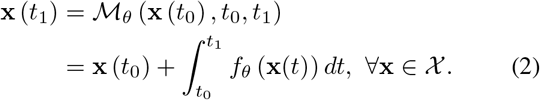

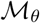 is differentiable with respect to *θ*. One may optimize the diffeomorphic flow to morph an initial surface at *t*_0_ towards a new surface at *t*_1_ that matches any target shape if the model has adequate capacity and the target shape is captured with enough data.

Given these considerations, we used a neural network-based ODE, or neural ODE [26], to model and optimize a diffeomorphic flow to morph a surface template to match the boundary points of the stomach. After exploring it with 2-D test examples, we used this strategy to model the rat stomach observed with MRI.

### B. 2-D test examples

To explore its utility, we first tested the neural ODE model with synthetic test examples with the same architecture and setting as for experimental data, except that the input coordinates were 2-D instead of 3-D. In the example shown in Fig. 1, an ecliptic template (blue) was deformed to match an iso-surface represented with a point cloud (red). When the neural ODE was optimized, it formed a nonlinear (curved) path to move each point on the template towards its best-matched target point (Fig. 1.b). The template as a whole was deformed progressively towards an optimal fit with all target points (Fig. 1.b). In contrast, when the same network used for the neural ODE was instead optimized to implement a discrete and nonlinear registration, the resulting contour failed to capture the expected shape of the iso-surface (Fig. 1.c). More test examples are shown in Fig. 2. The performance of deforming a template contour to match a target shape was high, regardless of whether the template began with a non-informative circle or a prior shape similar to the target, or whether the points sampled from the surface were dense or sparse.

**Fig. 1.**
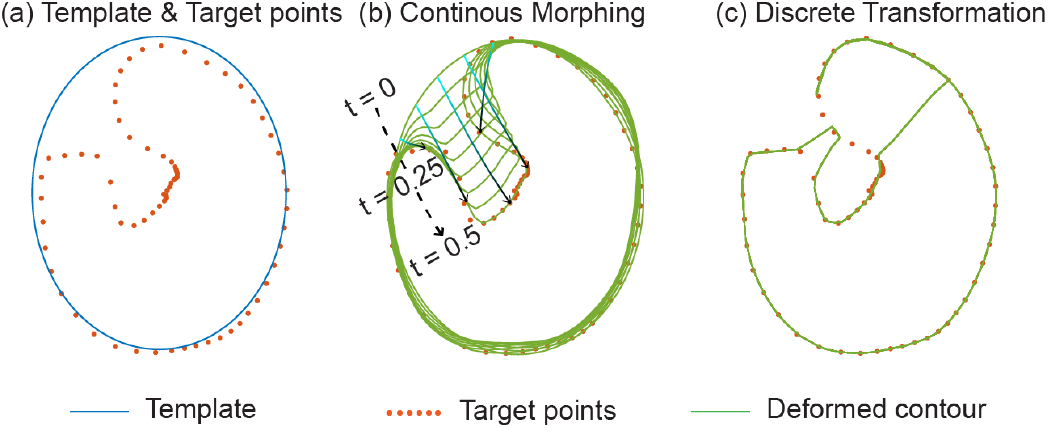
Continuous morphing vs. discrete transformation. (a) shows an example problem of aligning a 2-D template (a blue contour) to a set of target points (red). (b) shows the result of using a diffeomorphic flow to morph the template towards a progressive fit with the target points. A neural ODE was optimized and solved to continuously drive the template from its original shape at t=0 towards its final shape at t=0.5. The shapes at several time points between t=0 and t=0.5 are shown (green) to illustrate the process of morphing. Five example locations on the template evolve along unique trajectories (cyan to black encodes increasing time) towards the target points. For comparison, (c) shows the discrete mapping that transforms the template in an attempt to match the target points. The discrete mapping uses the same neural network architecture as the neural ODE model used in (b). The discrete mapping does not give rise to a continuous contour (green), while leaving some target points unmatched or mismatched.

**Fig. 2.**
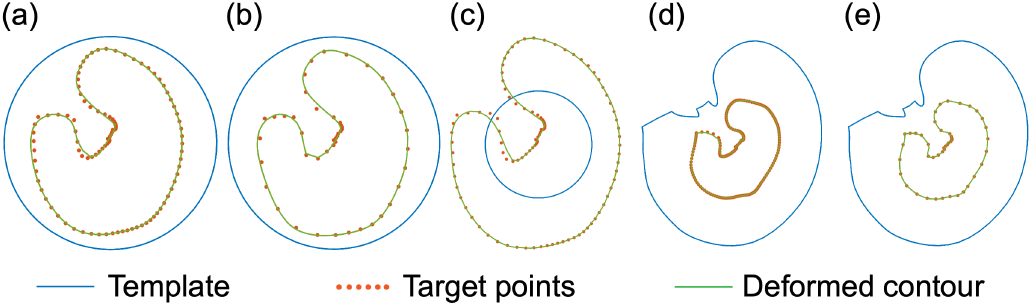
2-D test examples. In each example, a template (as a blue contour) was morphed to a new shape (green) that matches the target points (red). In (a) through (c), the template was a circle without any informative prior about the target points. In (d) and (c), the template started with a similar shape as the target points. In these examples, the target points are denser in (a) and (c) relative to (b), and denser (d) relative to (e). The target points cover a larger (c), smaller (d & e), or comparable (a & b) scale relative to the template. Regardless of these different conditions, the neural ODE can be optimized to morph the template to match the target points accurately.

### C. 3-D surface of a stomach

For a real stomach, we deformed a source surface to create a new surface that matched a target iso-surface surrounding the stomach as observed with MRI. Here, we used a mesh as the topological representation of a continuous surface. It was defined as a set 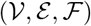 of vertices 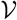, edges 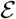, and faces 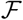. This mesh was based on a generic scaffold of the rat stomach (Section II-C.1). The target iso-surface was extracted from the image segmentation of the stomach (Section II-C.2) for capturing both the anatomy and the motility. The problem was thus defined as the optimization of the model for morphing the source surface to fit the target iso-surface (Section II-C.3, Section II-C.4, II-C.5, and II-C.6). During morphing, we kept the defined 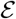 and 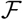 while updating 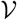.

#### 1) Surface template as the source

A shell was extracted from a digital 3-D rat stomach built from the micro-CT images from the 11 rat stomachs [22]. This shell was a surface mesh representative of the generic anatomy of the rat stomach. It was closed after separating the stomach from the esophagus and the duodenum and then closing the corresponding openings. It included 4,664 vertices and 9,324 triangular faces. We further refined this mesh by adding one vertex to each edge and dividing each face into four using [36]. The refined scaffold included 18,650 vertices and 37,296 faces and defined the source surface used in the subsequent analyses. Hereafter, we refer to this shell as the generic scaffold.

#### 2) Iso-surface of the 3-D volume as the target

An iso-surface was created to delineate the boundary of gastric lumen, which was contrast enhanced in MRI. Two types of iso-surfaces were of interest. One captured the time-averaged shape of the stomach (anatomy). The other captured the time-resolved shape of the stomach relevant to the dynamic movement of the stomach wall (motility).

The iso-surface was extracted from the 3-D gastric lumen segmented from MRI. Detailed information about MRI acquisition can be found in Section. III. After reconstruction, MRI data were resampled to 128 × 128 × 64 voxels of 0.5 mm isotropic resolution with trilinear interpolation. The stomach and the intestines, showing much higher intensity than elsewhere, were segmented using a 3-D U-net [37], [38]. The 3D U-net was trained with manually annotated MRI labels from 48 rats and validated with data from six held-out rats as described in our earlier work [39]. The labels assigned each voxel to the stomach, intestines, or background. The trained model was applied to MRI images from 10 rats used in this paper. The segmentation results were further confirmed by visual inspection as well as a conventional method based on intensity thresholding [40]. We used an iso-surface extraction method with a kernel size of three. When the segmentation was averaged over time, the extracted iso-surface represented a static stomach. When the iso-surface were extracted from the segmentation at each time, they represented a moving stomach.

The generic scaffold was spatially aligned to the target iso-surface through three stages (Fig. 3): 1) a linear affine transformation, 2) a neural ODE optimized to morph the scaffold to match the static iso-surface, and 3) another neural ODE model optimized to further morph the scaffold to match the dynamic iso-surface observed at each time point. The first and the second steps established a scaffold that captured the individualized anatomy. The third step captured the movement of the stomach for subsequent motility analyses.

**Fig. 3.**
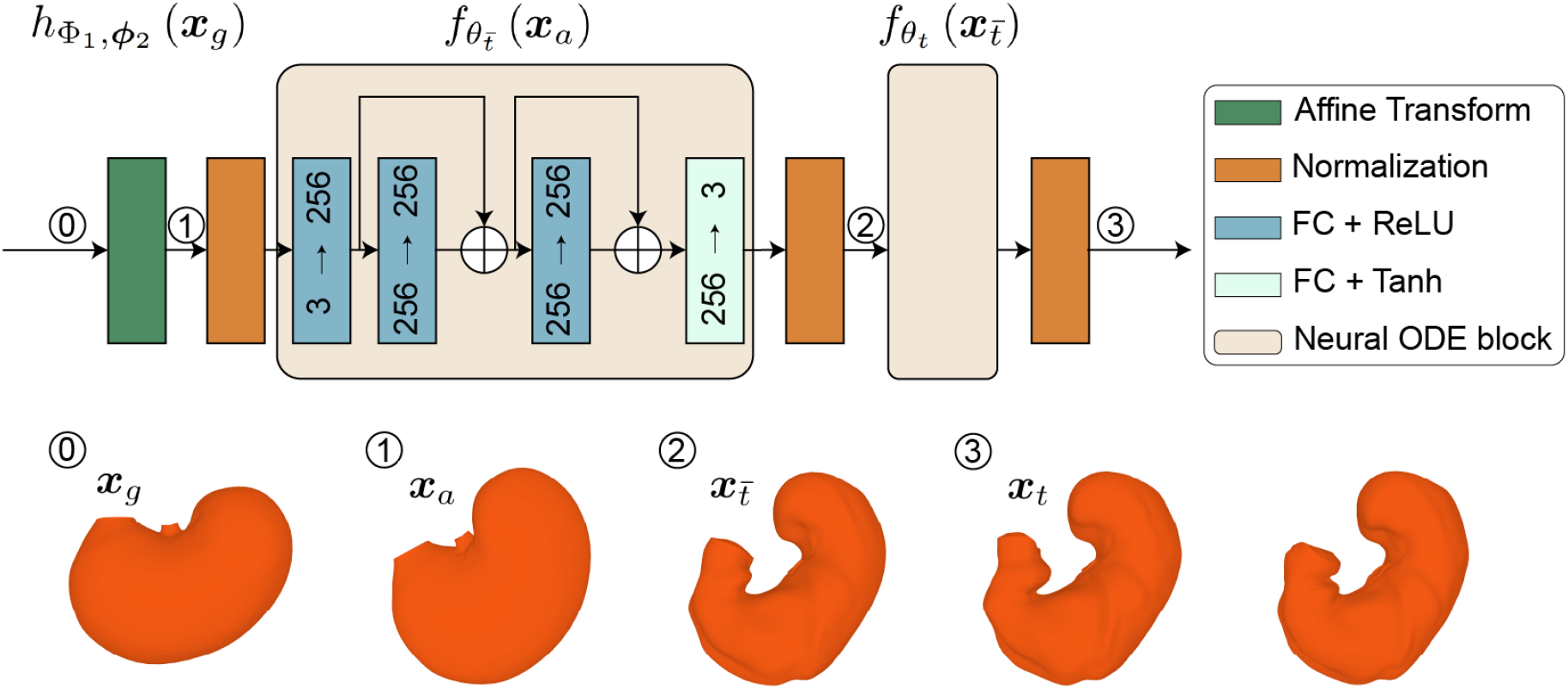
Model architecture. The top shows the architecture of the model used to align the generic scaffold to a moving iso-surface derived from MRI. The model implements a computational graph that includes three sequential stages. The first stage is a linear affine transformation 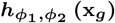 for initial alignment. The second stage is a nonlinear neural ODE block 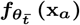 for alignment to the time-averaged MRI. The neural ODE block consists of a neural network with multiple layers as shown inside the block. The third stage is another neural ODE block 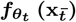 for alignment to the time-resolved MRI data. It uses the same network architecture as the neural ODE in the second stage. The bottom shows the scaffold in its original shape x_*g*_, following the affine transform x_*a*_, after being morphed to the static stomach 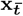, and further to a moving stomach observed at each individual time x_*t*_.

#### 3) Chamfer distance between source and target

The model was optimized to move the source to the target by minimizing the source-to-target distance. This distance was calculated as the Chamfer distance between points on the deformed surface and those in the target iso-surface. It is defined as *L_c_* in

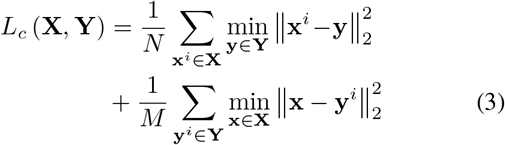

where *N* is the total number of sampled points x from the deformed source surface X whereas *M* is the total number of points y in the target iso-surface Y. Note that *N* is not necessarily equal to *M*. We further applied a regularizer to penalize the edge 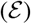 length [41], [42] as in

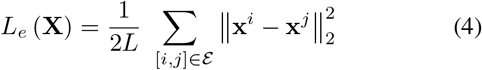

where *L* is the total number of edges in 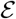. The Chamfer distance and the edge length regularizer were calculated using PyTorch3D [43].

#### 4) Affine tranform

In the first stage of aligning the source surface to the target iso-surface, we applied a linear affine transformation (Fig. 3, stage (0) to stage (1)). This step rescaled, translated, sheared, and rotated the generic scaffold X_*g*_ to roughly align itself with the static iso-surface, as well as two fiducial l andmarks (esophageal a nd pyloric openings). The linear affine transformation is defined as

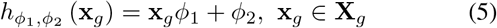

with *ϕ*_1_ ∈ ℝ^3×3^ and *ϕ*_2_ ∈ ℝ^1×3^. It was optimized to minimize the Chamfer distance *L_c_* between the affine-transformed scaffold 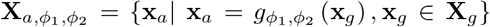 and the target iso-surface 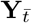 as expressed by

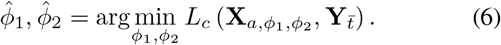

We implemented the algorithm using Python 3.6. To iteratively optimize the affine transform from a coarser to a finer level, we randomly sampled the vertices on the scaffold as x for calculating *L_c_*, starting with 2,000 and increasing it by 1,000 every 400 iterations. The Adam optimizer [44], as implemented in PyTorch, was used with an initial learning rate of 10^−3^, decayed by ten every 1,000 iterations for up to 4,000 iterations. The scaffold following the optimized affine transform was denoted as 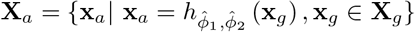.

#### 5) Morphing to a static stomach with Neural ODE

Following the linear affine transformation, the scaffold was morphed to match the static iso-surface through a neural ODE model. The model architecture is illustrated in Fig. 4. It used four fully-connected layers. The input channel was three, representing the 3-D coordinate. The dimension of the hidden layers was 256. This hyperparameter determined the capacity of the model and the complexity of the diffeomorphic flow that the model could support. The rectified linear unit (ReLU) was used after all hidden layers. The hyperbolic tangent (Tanh) function was used at the output layer, enabling both negative and positive flows [45]. As a whole, the neural ODE model described a nonlinear and Lipschitz-continuous function *f_θ_* (x(*t*)). We centered and normalized all input coordinates such that they were all within a unitary sphere at the origin before applying the neural ODE and then re-centered and re-scaled the deformed coordinates after applying the neural ODE.

**Fig. 4.**
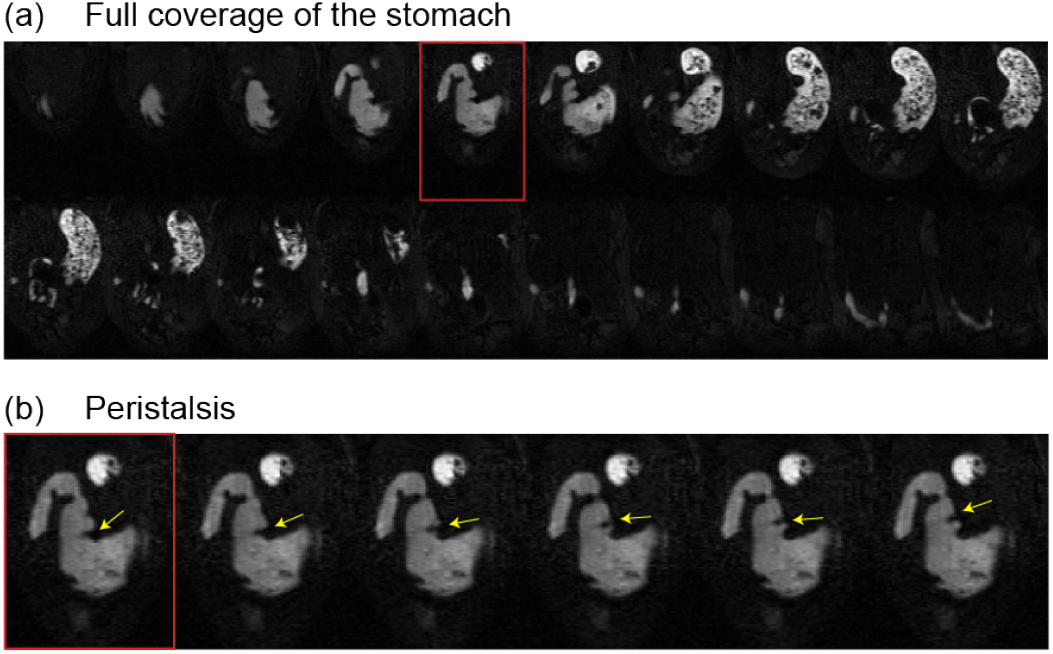
Representative gastric MRI data enclosing the intraluminal volume as well as capturing gastric motility. (a) shows 20 MRI slices that formed a 3-D volumetric description of the stomach at one fixed time point. The luminal content in the GI tract is bright in contrast to surrounding tissues or other viscera (as the dark background). For the slice (highlighted in red box), (b) shows 10 continuous scans separated by 2-3 s. The contraction is most visible in the antrum. The yellow arrow highlights a strong contraction (inward bending) that moves its position over time.

Mathematically, we described the morphing at this stage using neural ODE as 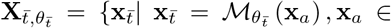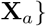. Here, for clarity, we omitted *t*_0_ and *t*_1_ from the input of the mapping 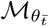 since we used them as two fixed hyperparameters in our implementation. The optimization of the model parameters 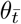 was expressed in

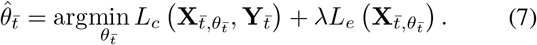

The optimized neural ODE resulted in the static scaffold denoted as 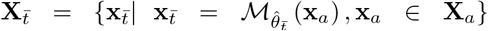, resulting from morphing the affine-transformed scaffold to match the time-averaged iso-surface 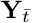. Here, we added the edge regularizer *L_e_* to enforce the regularity of the edges. Hyperparameter *λ* determined the relative importance of the regularizer.

The dopri5 solver implemented in PyTorch [26] was used. The absolute and relative tolerance was set to 10^−6^ to balance the computational time and accuracy. The integration time *t*_0_ and *t*_1_ were set as 0 and 0.2, respectively. To optimize the neural ODE, we attempted to morph the scaffold for the target points to fall onto the faces of the morphed scaffold, rather on aligning the vertices to the target points. For this sake, we uniformly sampled 10k points from the faces on the scaffold for calculating the Chamfer distance *L_c_*. The hyperparameter *λ* was set to be 1. We used the Adam optimizer for up to 1600 iterations with an initial learning rate of 10^−3^. The learning rate was decayed by ten every 400 iterations until reaching its minimum value of 10^−6^.

#### 6) Morphing to a moving stomach with Neural ODE

Another neural ODE model was utilized to morph a time-averaged scaffold 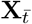 to capture the time-evolving motility. The neural network architecture was kept identical to the second stage. In this stage, we defined the output of the neural ODE as 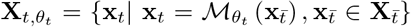. Subscript *t* referred to a specific time at which the target iso-surface was extracted. To optimize the model parameters *θ_t_*, we used the equation in

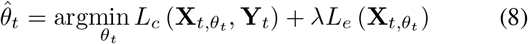

where Y_*t*_ was the time-specific iso-surface at time *t*. Once optimized, the morphed output is denoted as 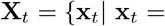 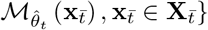.

The absolute and relative tolerance was decreased to 10^−3^ to reduce the computational time further. The hyperparameter *λ* was set to 0.1 for capturing a larger local deformation. We used the Adam optimizer for up to 2400 iterations with an initial learning rate of 10^−3^. The learning rate was decayed by ten if the loss was not decreased for 100 iterations. The optimization was stopped earlier if the loss was not decreased for 50 iterations after the learning rate reached its minimum of 10^−6^. Except for these differences, we used the same setting as in Section II-C.5.

### D. Characterization of the gastric motility

Relative to the time-averaged scaffold, we evaluated the displacement of every surface location at each time point, providing a location-specific time series of gastric motor events. The amplitude of the displacement was calculated as the 3-D Euclidean distance relative to the time-averaged reference position, whereas the sign was determined as positive or negative if the displacement was in a direction parallel or anti-parallel to the normal direction of each location. The time series of displacement was evaluated to identify the frequency, amplitude, and coordination of contractions observed at each location. After these features were analyzed in each subject, they were further averaged across subjects to report group-level motility characteristics.

#### 1) Frequency

To characterize the frequency of contraction, the time series of displacement was resampled at 0.5 Hz. Morlet wavelet transform [46] was applied to the resampled time series to represent the signal power as a function of time and frequency. Averaging this time-frequency representation over time resulted in a power spectrum, indicative of the frequency feature at each location on the stomach.

We evaluated how often the motility time series had its peak frequency between 3 and 7 cycles per minute (cpm), which is the frequency range of gastric slow wave initiated and paced by the interstitial cell of cajal (ICC) [47]. We counted the number of sessions that oscillating contractions occurred at the slow wave frequency relative to the total number of sessions. The percentage of the counted occurrence was used to indicate the likelihood that an oscillating contraction at the slow-wave frequency was observable at each location on the stomach.

We also evaluated the extent to which the oscillating contraction between 3 and 7 cpm accounted for the deformation observed at each location. This metric, termed as rhythmicity, was calculated as the ratio (in terms of percentage) between the power at the slow-wave dominant frequency (±1 cpm) and the power summed over all frequencies (from 0.1 to 12 cpm).

#### 2) Amplitude

We further band-pass filtered the motility time series around the dominant frequency (±1 cpm) to estimate the slow wave of contractions. The peak-to-peak amplitude was then extracted and averaged over time, labeled as the time-average amplitude of phasic contraction or contractile amplitude.

#### 3) Coordination

To identify the stomach-wide coordination of peristaltic contraction, we quantified the time-evolving phase at each location by applying the Hilbert transform to the band-pass filtered motility signal (the passing band was ±1 cpm around the dominant frequency). The difference in the instantaneous phases (relative phases) was calculated between each location on the stomach and a reference location on the antrum. Note that this relative phase is expected to be relatively stable, if the contraction at a location is coordinated with that at the reference location due to the propagation of peristaltic wave. In contrast, if the two locations are not coordinated, then the relative phases would vary over time and tend to average to zero. Therefore, to examine the level of coordination, we averaged the relative phase over time, resulting in a relative phase in a range from −*π* to *π* (in radians).

### E. Characterization of the structure-function relationship

From recent studies [1], [29], structural information about the gastric musculature in rat has been collected [1] and registered to the same scaffold [29], onto which we mapped the motility. Such co-registered structural (musculature) and functional (motility) features allowed us to evaluate their relationship. The structural features included the thickness of the circular muscle (CM) layer and the longitudinal muscle (LM) layer. The functional features pertained to the frequency, amplitude, and coordination of muscle contractions. Upon initial exploration and visual inspection, the correlations between the contractile amplitude and the thickness of CM, LM, or both appeared to fall into two distinct ranges, suggesting that the stomach may be separated into two functional regions with distinct structure-function relationships (between muscle thickness and motility). To delineate the two regions, we concatenated the structural and functional features after standardizing each feature and then performed k-means clustering of the concatenated features with k=2. The resulting two clusters grouped all locations on the stomach into two non-overlapping regions, which were referred to as the proximal and distal clusters based on their distributions. Separately for each cluster, we tested whether the thickness of CM *x*_1_ and the thickness of LM *x*_2_ could explain the contractile amplitude *y* through a linear regression model: *y* = *α*_1_*x*_1_ + *α*_2_*x*_2_ + *ϵ*. The regression coefficients were further used to evaluate the differential contributions from the CM and LM (*α*_1_ vs. *α*_2_) to the amplitude of peristaltic contractions. An F-test was performed to test the overall significance of the model (the effects of CM and LM on the gastric motility).

## III. EXPERIMENTS

Ten Sprague-Dawley rats (male, 250-350g, Envigo, Indianapolis, Indiana) were used in this study. The animals were housed in a room with a controlled room temperature of 68 - 79°F, relative humidity from 30% to 70%, and light on from 6 am to 6 pm. All animal procedures followed a protocol approved by the Unit for Laboratory Animal Medicine and the Institutional Animal Care & Use Committee at the University of Michigan. The animals first underwent a diet training procedure [12], such that each animal was able to voluntarily consume a 5-g meal (DietGel Recovery, ClearH2O, ME, USA) labeled with Gd-DTPA (SKU 381667, Sigma Aldrich, St. Louis, USA) 15 to 30 minutes prior to MRI. The animal was initially anesthetized with 4% isoflurane mixed with oxygen for up to 5 min and maintained with 2.5% isoflurane until it was set up on the animal holder in a prone position and ready for MRI scanning. Before the image acquisition, the animal was further sedated with a bolus injection of dexmedetomidine (0.05 mg/ml, Zoetis, NJ, USA) at 0.01 mg/ml, 0.0125 mg/kg. Afterwards, the animal was kept stable with a combination of 0.5% isoflurane and 0.01 mg/ml, 0.025 mg/kg/h dexmedetomidine by continuous infusion. Throughout the experiment, the animals were stable, with respiration rates of 30 - 60 cpm.

GI MRI [28] was performed using a 7-tesla small-animal MRI system (Varian, Agilent Technologies, California, USA) with a volume transmit and receive 1H RF coil with a 60mm inner diameter. Images were taken with a multi-slice gradient echo sequence with echo time = 1.44 ms, repetition time = 10.50 ms, flip angle = 25°, the number of slices = 20, field of view = 64 mm × 42 mm, matrix size = 128 × 84, slice thickness = 1.5 mm. The slices were prescribed in the oblique direction in parallel with the long axis of the stomach, with phase-encoding in the left-right direction and frequency-encoding in the head-tail direction. The acquisition was gated by the animal’s respiratory trace to mitigate the effect of respiratory motion. A spatially non-selective saturation pulse was applied before image acquisition to enhance the T1-weighted contrast. The entire acquisition was achieved with multiple sessions and lasted around 4 h per animal.

## IV. RESULTS

In this study, we morphed the generic scaffold to match the MRI data acquired in ten healthy rats. The iso-surface (boundary points) of the intraluminal volume were used as the target for generating static and dynamic surface representations of the stomach from the generic scaffold. The gastric anatomy and motility were characterized and mapped for each individual and then summarized at the group level. The group-level motility and structural maps was then co-registered and compared to address the relationship between muscle thickness and gastric motility.

### A. Gastric MRI

After each rat consumed 5-g Gd-labeled diet-gel, the ingested diet was visible with T1-weighted MRI. Fig. 4 shows the MRI images for 20 slices sampled every 2 to 3 s from a representative rat. Voxels in the lumen showed higher contrast against those outside. The boundary of the luminal content approximated the inner surface of the stomach (Fig. 4.a). The change of this boundary reflected the movement of the stomach wall. Fig. 4.b highlights the dynamic changes on one single slice. Deformations of the gastric boundary indicate contractions in the distal stomach (primarily the antrum). Its peaks and troughs moved over time from the proximal towards the distal part of the organ, suggesting a traveling wave. In this example, the wave occurred with a mean frequency of 5.390± 0.996 cpm. In the antrum, the luminal contour showed greater inward bending than outward bending (Fig. 4.b), suggesting that the contraction was phasic and possibly occlusive. The contraction was stronger at the lesser curvature than at the greater curvature.

### B. Morphing with neural ODE

The generic scaffold was affine-transformed and then morphed to match the time-averaged shape of the stomach by an optimized neural ODE model (Section II-C.5). The model parameters were optimized iteratively and tended to converge after 200 iterations and reached the optimum up to 1,200 iterations (Fig. 5.a). After it was optimized, solving the neural ODE was a continuous process that unrolled itself over time to progressively transform every point on the scaffold until the morphed scaffold matched the target iso-surface derived from MRI, as illustrated in Fig. 5.b. This morphing procedure could generate a static scaffold representing the stomach of every rat (Fig. 6). Similarly, a separate neural ODE model was optimized to morph the static scaffold to match the MRI data observed at each time point. By visual inspection, dynamic MRI data (Fig. 7.c) and its corresponding scaffold obtained by the neural ODE (Fig. 7.d) closely matched each other in terms of the morphological features and their changes over time. The dynamic scaffold demonstrates a propagating wave of muscle contraction – a hallmark feature of gastric motility under postprandial conditions.

**Fig. 5.**
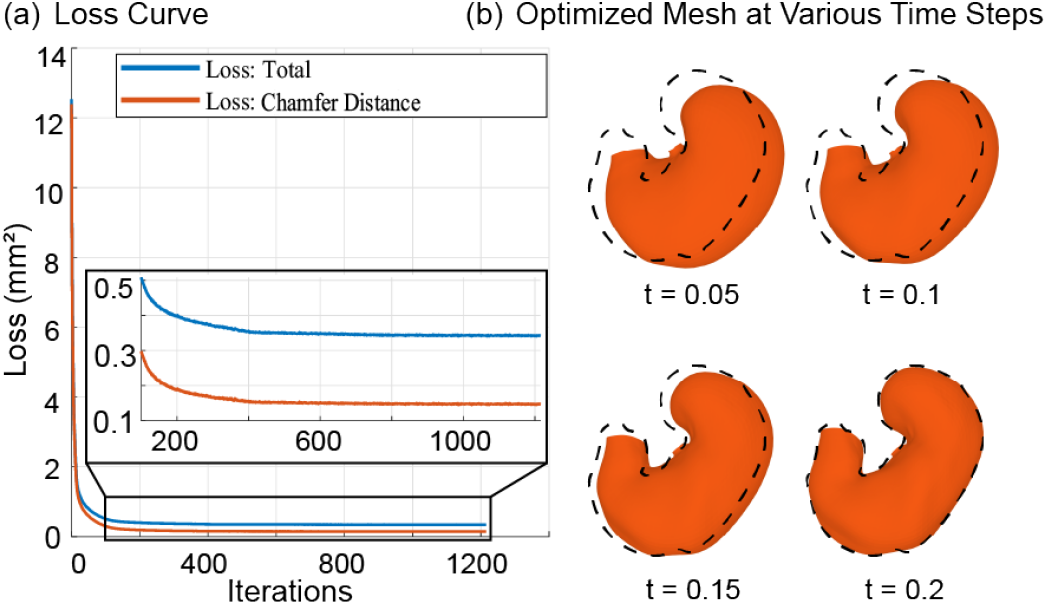
Optimization of the neural ODE. The iterative optimization of the neural ODE is shown in (a), representing the loss obtained at each iterative step. The Chamfer distance is shown as the red curve. The total loss, including the Chamfer distance and the edge length regularization, is shown as the blue curve. The insert figure in (a) highlights where convergence emerges during the iterative optimization. (b) illustrates the process of solving the optimized neural ODE. It shows the resulting scaffolds morphed until several time points t= 0.005, 0.1, 0.15, 0.2. The 3-D shape of the scaffold progressively approaches the contour of the target iso-surface (as the dashed line).

**Fig. 6.**
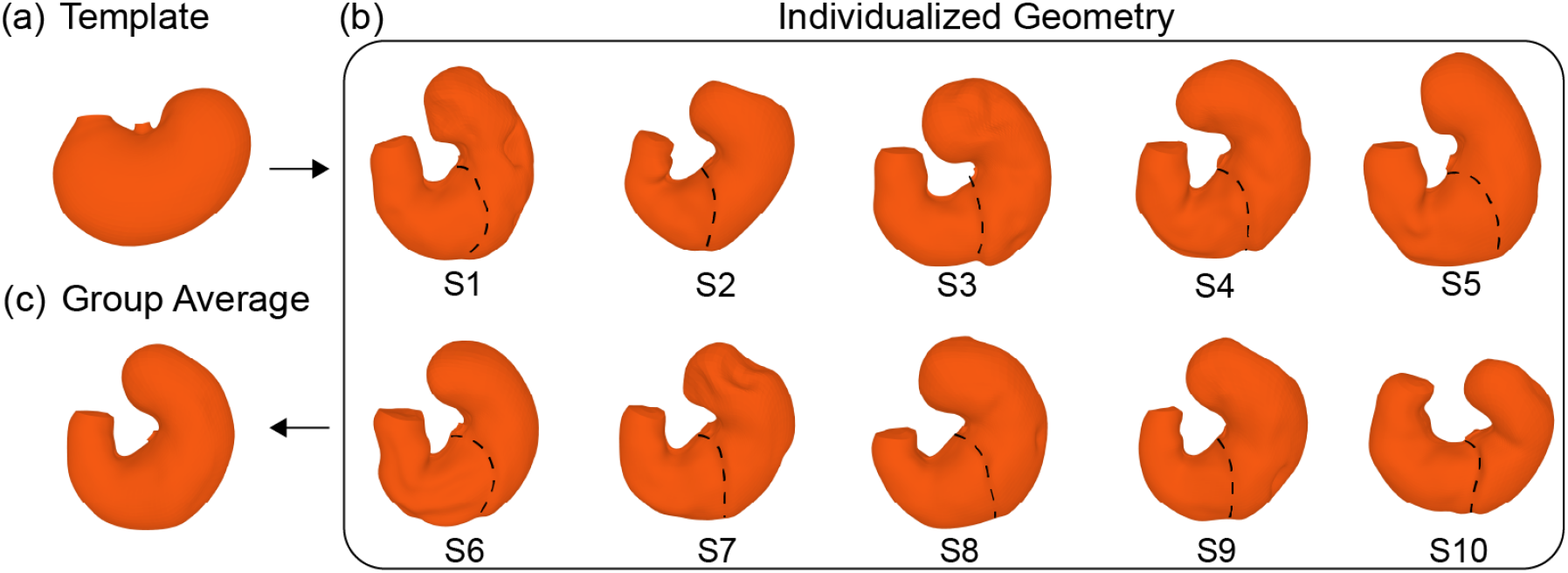
The geometry of the stomach at both individual and group levels. (a) shows the generic scaffold. (b) shows the individualized scaffolds after aligning the generic scaffold to MRI data for ten subjects (S1 through S10). In each subject, the limiting ridge is shown as the dashed line. (c) shows the group-level scaffold obtained by averaging the individualized scaffolds across all subjects.

**Fig. 7.**
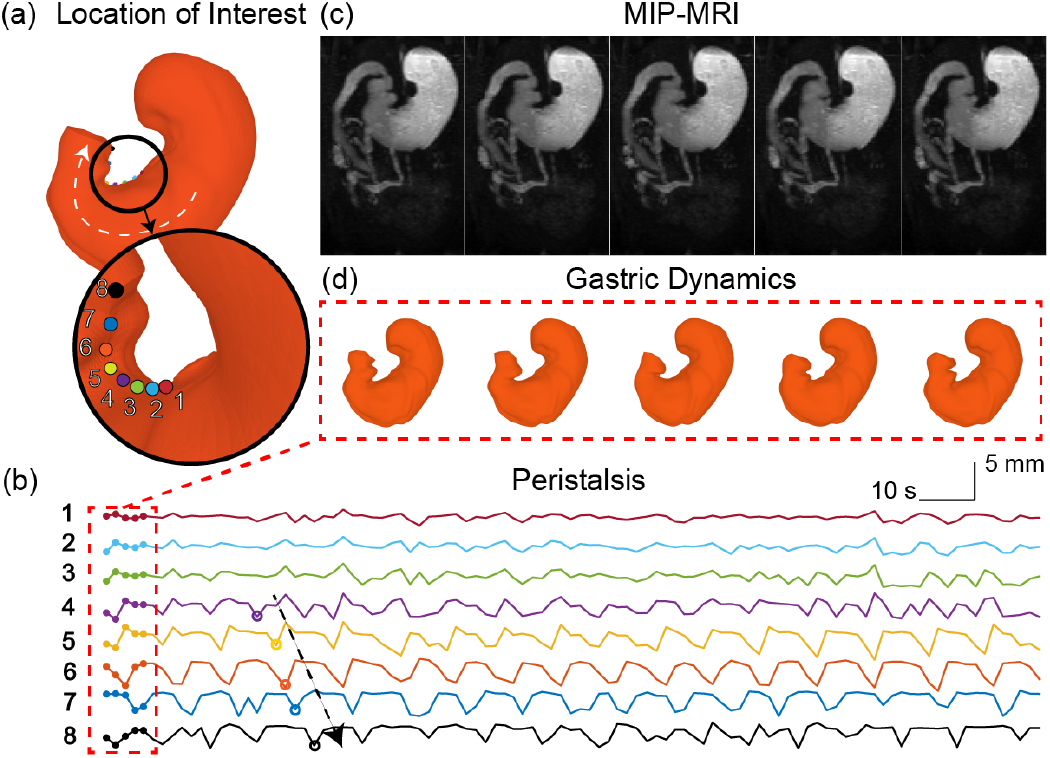
Gastric motor events. On the stomach model for a single subject, (a) shows eight locations of interest (numbered and color-coded) along the lesser curvature. (b) shows how each of these locations moves over time. The dashed arrow indicates a contraction (sharp trough) moving from the forth location towards the eighth location. At each of the five time points (in the red dashed box), the maximal intensity projection (MIP) of the MRI data is shown in (c) and the corresponding scaffold is shown in (d). The scaffold captures the surface movement consistent with MRI.

For the static scaffold, its error in matching with the static stomach in each rat was quantified as the Chamfer distance between the points on the scaffold and the points on the target iso-surface derived from the time-averaged MRI data. For all rats, this distance was 0.127 ± 0.020 mm (mean ± standard deviation), suggesting that the error of morphing was less than the voxel size (0.5 mm 0.5 mm 0.5 mm).

The morphed scaffolds preserved their integrity and were physically reasonable representations of the gastric surface. This benefits from the diffeomorphic flow enforced by the neural ODE. To verify whether the diffeomorphic prior was successfully enforced, we estimated the percentage of self-intersections of the morphed scaffolds. In our implementation in ten rats, we found no self-intersections in the surface of the static stomach, while the Chamfer distance remained small 0.197 ± 0.023 mm. The dynamic surface of the moving stomach showed a very low percentage of intersection 0.028%±0.170%. This indicates the implemented neural ODE successfully enforced the integrity of morphed scaffolds.

### C. Individual level

The static scaffold captured the individualized gastric anatomy and allowed comparison between rats. Fig. 6.b shows the static scaffold representing the time-averaged shape for each of the ten rats studied. Following a 5-g meal, all rats showed a common morphological feature for their stomachs. The individualized scaffold showed an oblique U-shape with the fundus and the antrum being much more elongated than the generic scaffold in Fig. 6.a. In every rat, the limiting ridge was observable as deep grooves on the ventral or dorsal surface (Fig. 6.b). Across different rats, the limiting ridge was found to start at roughly similar locations on the lesser curvature relative to the esophageal opening but end at more varying locations on the greater curvature (Fig. 6.b). This observation agrees with [1], [23]. Furthermore, individual variations could also be noted in terms of the relative sizes of the compartments separated by the limiting ridge, and the angle and width of the U shape (Fig. 6.b).

The dynamic scaffold provided rich spatiotemporal data about gastric motility and its patterns in each rat. The motility was observable on the scaffold. In a representative rat, we chose eight locations of interest along the lesser curvature (Fig. 7.a) and plotted their motion over time as a time series of motor events at each of these locations (Fig. 7.b). Muscle contractions and relaxations correspond to the troughs and peaks, respectively. As shown in Fig. 7.b, the motion time series showed different characteristics at different locations. A stronger motion was observed at more distal locations (closer to the pyloric opening) than at more proximal locations (closer to the esophageal opening). The motion was more periodic in the antrum (the last five rows in Fig. 7.b) than in the corpus (the first three rows in Fig. 7.b). The motion time series demonstrates asymmetric contraction-relaxation patterns at some locations, showing a stronger contraction over a shorter period followed by a weaker relaxation over a relatively longer period. A progressive phase delay was observable for five of the eight locations, suggesting a wave propagating from the proximal antrum to the distal antrum. The frequency and velocity of this wave were 6.105 ± 0.741 cpm, and 0.645 ± 0.129 mm/s, respectively. This measurement is consistent with the result from volumetric analysis reported in a previous study [12]. In the proximal stomach, oscillatory contractions did not show a clear propagation pattern. The relative phase of contraction between locations was disperse and varying, with a time average approaching zero. These findings generally agree with the quantification of gastric motility with the ex-vivo preparation by [6], [48].

### D. Group level

For every rat, the scaffold used a common topology such that the anatomical and motility characteristics could be readily averaged and compared across rats. Averaging the individualized scaffold across all rats generated a representative scaffold of the rat stomach (Fig. 6.b) following a 5-g meal. This group-level scaffold averaged out individual variations and preserved only the common morphological features.

Using this group-level scaffold, we chose five locations of interest (Fig. 8.a) and characterized the group-level motility at each of these locations. The group-averaged power spectral density showed different frequency characteristics at different locations (Fig. 8.a). The fundus (1 & 2) showed high-frequency and low-amplitude motion without a well-defined characteristic frequency. The corpus (3) and the antrum (4 & 5) showed motion with a progressively larger amplitude in a narrow range frequency from 3 to 7 cpm. A characteristic frequency was more pronounced in the antrum than elsewhere.

**Fig. 8.**
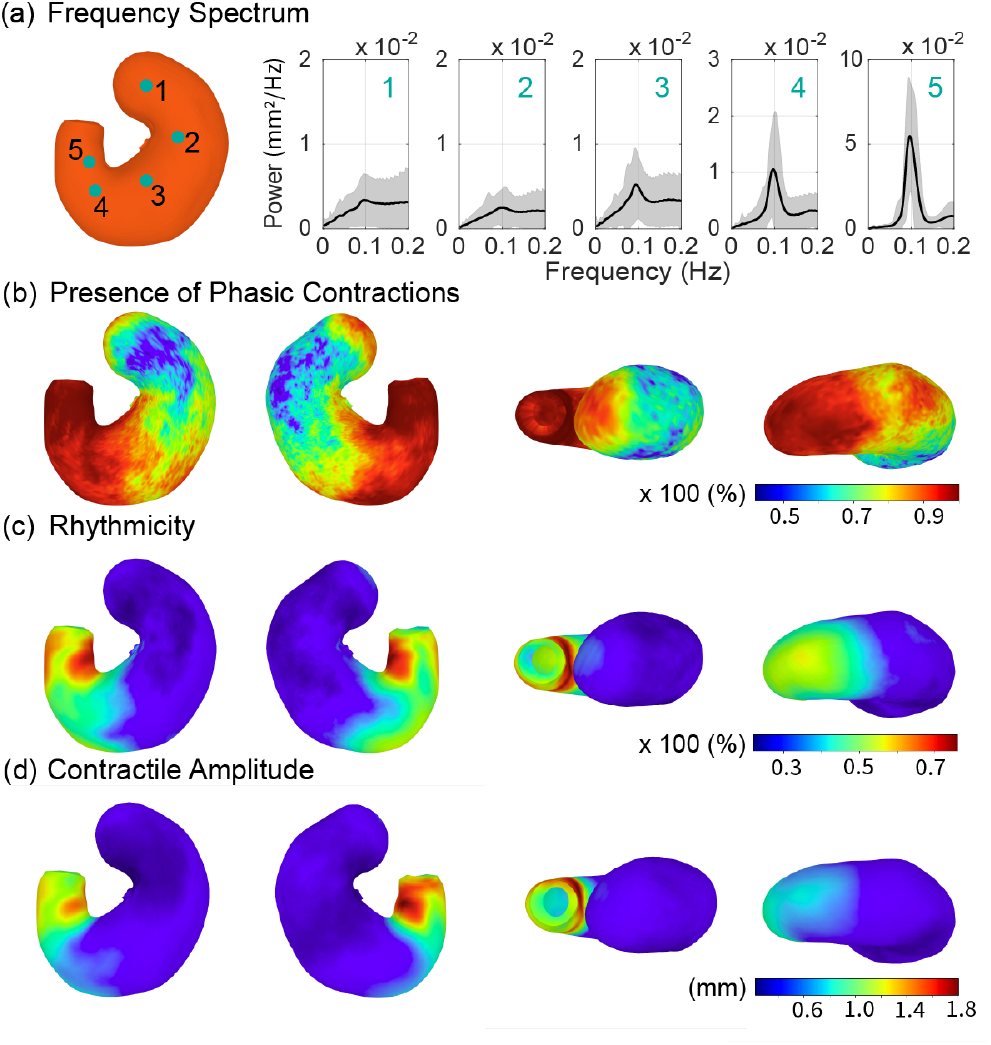
Motility characteristics. (a) shows the power spectral density (PSD) at five representative locations at the proximal fundus (1), distal fundus (2), proximal corpus (3), distal corpus (4), and antrum (5). The PSD is averaged across all sessions and subjects. The black curve is the group average. The grey zone shows the standard deviation about the average. (b) shows the percentage by which the motor events at each location exhibit a spectral peak between 3 and 7 cpm. (c) shows the relative contribution of periodic phasic contraction to the motor events observed at each location. (d) shows the peak-to-peak amplitude of phasic contraction at each location. The maps shown in panels (b) through (d) are viewed from the ventral, dorsal, top, or bottom perspective (arranged from left to right).

We further summarized the motility attributes and presented the results as a set of functional maps visualized on the group-level scaffold. For example, the percentage by which oscillatory motion between 3 and 7 cpm was present at each location was used to map where phasic contraction occurred in the stomach (Fig. 8.b). In general, phasic contractions were increasingly observable from the proximal to the distal stomach. The occurrence of phasic contraction showed a spatial gradient on the ventral and dorsal surfaces. Two regions stood out as exceptions. The fundic cap showed more phasic contractions than other fundic regions. The distal fundus on the greater curvature showed more phasic contractions than its surrounding regions. Besides, the “rhythmicity” map (as defined in Section II-D) showed that the antrum was most dominated by phasic contraction but with observable distinctions along the circumferential direction (Fig. 8.c). The lesser curvature and the greater curvature showed stronger rhythmicity than other ventral or dorsal locations in between (Fig. 8.c). The amplitude of phasic contraction showed a similar distribution (Fig. 8.d). It was further clear that phasic contraction was stronger on the lesser curvature than on the greater curvature (Fig. 8.d). It should be noted that the regional distinctions revealed by these functional maps do not follow the mucosal anatomical landmark [1] represented by the limiting ridge.

We further evaluated how phasic contractions coordinated between locations. We used each of the five locations of interest as a reference and calculated the relative phases. Fig. 9 showed a group-averaged map. When the reference point was located in the antrum, the relative phase spanned the full range from *π* to *π* and revealed a wave-like spatial profile (Fig. 9.a & b), suggesting that a robust peristaltic wave passed through the reference point. This peristaltic wave started from the distal corpus and ended at the distal antrum, spanning roughly two wavelengths. However, when the reference point was in the proximal corpus (Fig. 9.c) or fundus (Fig. 9.d & e), the relative phase was mostly around zero, suggesting the absence of a peristaltic wave propagating through the reference point.

**Fig. 9.**
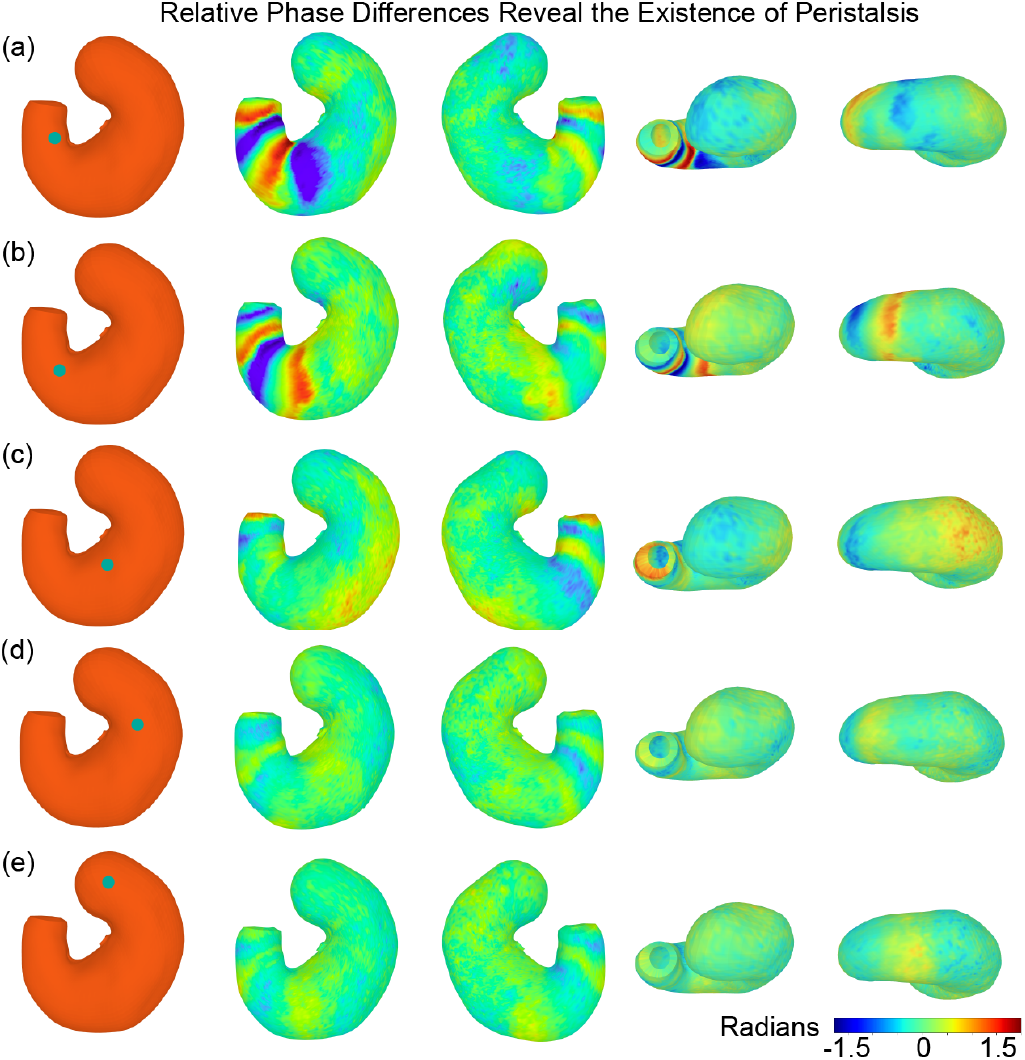
Coordination of gastric motility. Each of the five locations of interest served as the reference location to which the relative phase of contraction at every location on the stomach is visualized from *π* to *π*. The reference location is shown as the red dot in panels (a) through (e). The relative phase is visualized on the surface of the stomach as viewed from the ventral, dorsal, top, or bottom perspective (arranged from left to right) in each panel.

We further compared and integrated the information about muscle thickness [1] and gastric motility (reported herein) co-registered on the same scaffold of the rat stomach. The stomach was parcellated into two functional regions based on k-means (k=2) clustering of the combined structural and functional features available at each location of the stomach, including LM and CM thickness, the frequency, amplitude, and coordination of gastric motility. The two functional regions were referred to as the proximal cluster and the distal cluster, given their relative orientation and position, as in Fig. 10.c. The proximal cluster included the fundus and the proximal corpus, whereas the distal cluster spanned across the distal corpus, the antrum, and the pyloric sphincter. The boundary between the two clusters was drawn as a white curve. This boundary was more towards the distal stomach relative to the limiting ridge. Around this boundary, the thickness of both CM (Fig. 10.d) and LM (Fig. 10.e) showed a smooth gradient, instead of any notably sharp change. Relative to this boundary, the more proximal locations showed much lower contractile amplitudes (Fig. 10.f). The proximal and distal clusters clearly showed different relationships between the contractile amplitude and the CM or LM thickness. A much stronger correlation was observed for the distal cluster than the proximal cluster. Fig. 10.a and b illustrate the corresponding scatter plots, showing a “L-shaped” mixture of two correlational relationships. The CM and LM thickness could explain the contractile amplitude for both the distal and proximal clusters, through linear regression models, *y* = 0.43*x*_1_ + 0.18*x*_2_ and *y* = −0.53*x*_1_ −0.11*x*_2_, respectively. While both models were statistically significant (F-test, *p* < 0.001), the contractile amplitude depended more on the CM thickness than the LM thickness.

**Fig. 10.**
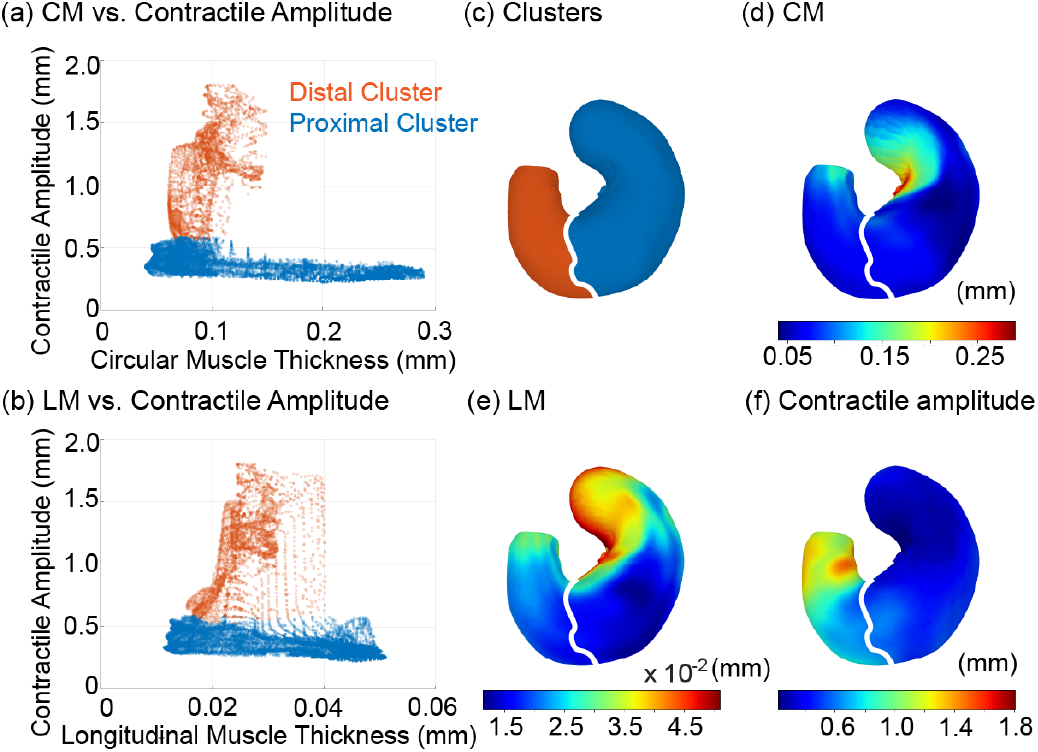
The relationship between muscle thickness and contractile amplitude. (a) shows a scatter plot of the relationship between the CM thickness and the contractile amplitude at locations in the proximal cluster (blue) or the distal cluster (red). (b) shows a similar scatter plot for the LM thickness. (c) shows the locations in the proximal cluster (blue) and the distal cluster (red) and their boundary (white). (d) and (e) show the distributions of the CM and LM thickness and the boundary that separates the proximal and distal cluster. (f) shows the distribution of the contractile amplitude alongside the boundary between the proximal and distal clusters. All maps are displayed on the scaffold representing the group-averaged stomach.

## V. DISCUSSION

The approach described here enables in-vivo and non-invasive characterization of gastric anatomy and motility through surface-based morphological analyses of gastric MR images in rats. Our approach has advantages over alternative methods that use invasive or ex-vivo procedures to measure muscular activities at limited locations. It can be used to identify various motor events throughout the surface of the stomach. It provides sub-millimeter spatial resolution and 2 to 3 s of temporal resolution. Importantly, our approach supports both individual and group-level analyses of gastric anatomy and motility. A deformable template is used to model the luminal surface and its movement at different times, conditions, and subjects. This allows one to compare the motility patterns within and across subjects, or potentially between healthy and diseased conditions for future work. It also allows one to co-register and compare structural and functional data and to further relate the MRI-observed motor events, such as gastric motility, to the underlying structural information, such as the musculature. The following discussions are extended from the perspectives of modeling, simulation, physiology, and clinical translation.

A common strategy for comparison across individual surfaces is to generate a mesh from individualized images and then align the generated meshes to a common template. Our approach uses a different strategy. That is to morph a fixed surface template to fit individualized images. The scaffold captures the generic shape of the rat stomach and bears an intrinsic shape prior of the stomach across different conditions and individuals. The correspondence between this scaffold and the individualized stomach is established through two stages of alignment, as in Section. II-C.4 and II-C.5. Both two stages use the time-averaged stomach as the target for alignment. This is because the dynamic stomach has its shape influenced by muscle contractions, whereas the generic scaffold reflects the ex-vivo stomach without any contraction. It is preferable to preclude the time-dependent effects of contractions on the alignment from a static template to a static stomach.

A key innovation of our approach is the use of neural ODE to capture gastric motility. The neural ODE learns a diffeomorphic flow to continuously morph the stomach. Such continuous morphing is physiologically plausible, because biomechanics ascribes the force-deformation relationship to a continuous, differentiable, and nonlinear process. With neural ODE models, we introduce this inductive bias into the deformation, ensuring that any two points on the surface would not collide with each other. Theoretically, this inductive bias also ensures that the surface after morphing remains a smooth surface (manifold), as long as the surface before morphing is smooth. In contrast, a discrete transformation may disrupt the manifoldness (see Fig. 1).

In a related study, [27] proposed a similar model with simultaneously learned shape embeddings through a PointNet [49]. They started with a non-informative spherical template and deformed it to match an arbitrary object with a shape-embedding guided diffeomorphic flow. In contrast, our study exploits the diffeomorphic flow as a physiologically plausible prior for registering a deformable organ – the stomach. For this purpose, we start with an anatomically realistic template of the stomach and morph it to match the boundary points of the intragastric content extracted from MRI. The template includes shape priors about the stomach, such as its curved “U” shape, pyloric and esophageal openings, greater and lesser curvatures, the asymmetry between the proximal and distal ends, and the symmetry between the ventral and dorsal sides. Therefore, it is not necessary to learn any shape embedding from the target surface. The neural ODE model alone can implement a gradual morphing process that begins with the surface template and ends up with an accurate surface representation of any static or moving stomach.

In our work, the neural ODE that morphs from a static stomach into a moving stomach is optimized separately for each time point of observation. This choice is conservative because it assumes each time point is an independent sample. Although this assumption does not compromise the accuracy of morphological characterization, it may be relaxed in future studies. The movement of the stomach typically exhibits regular temporal patterns. Such temporal or spectral features may be exploited to ease the optimization of the neural ODE. For example, it is likely feasible to use a single neural ODE as a dynamic system to model how the stomach moves over a period of time, similar to what was proposed in [50]. In our case and for future studies, such a dynamical system may be implemented as a second-order (or higher-order) ODE [51] to predict the movement of the stomach even beyond the period of observation, providing a generative model to simulate gastric motor events. One may use this model to interpolate missing samples to increase the temporal resolution or extrapolate future samples to reduce the scan time.

Our approach generates a smooth and moving mesh for the luminal surface of the stomach. As its topology is pre-defined and preserved by the template scaffold, this mesh sets up a moving boundary condition for a simulator of computational fluid dynamics. Given further information about pyloric opening, one can predict food transit to understand the relationship between gastric motility and emptying.

Similar analyses may be extended from the stomach to the entire GI tract. From MRI, the labeled food is also visible in the intestine. A multi-compartmental cylinder may be used as a template for the GI tract. This natural extension is expected to bring synergistic capabilities for addressing the coordination among different GI segments, such as the coordination of the antrum, pylorus, and duodenum. This template may be generated from the concatenation of multiple scaffolds, each representing one segment of the GI tract.

Our approach generates rich spatiotemporal data about gastric motility. This paper focuses on the methodology, instead of novel physiological findings. Our results demonstrate the feasibility of revealing high-resolution functional maps of gastric contractile activity in healthy rats. The obtained motility features include but are not limited to the amplitude, frequency, propagation, and coordination of contractile activity. In this study, such functional features in combination with structural features of the gastric musculature were used in parcellating the stomach into two structurally and functionally distinct regions. In future studies, more fine-grained anatomical and functional parcels may be attainable by including more anatomical features, such as the vasculature, enteric nervous system, and extrinsic neural innervation. It is interesting to note that the amplitude of the peristaltic contraction depends more on the CM thickness than the LM thickness. Such a differential effects from different muscle layers may be attributed to the alignment and organization of CM and LM muscle fibers relative to the direction of peristaltic propagation. One limitation in this study is that the muscle thickness was evaluated from the isolated tissue samples from the ex vivo stomach, whereas the gastric motility was evaluated in vivo with MRI and under post-prandial conditions. The muscle thickness is expected to vary with the volume of the intragastric ingesta. A more rigorous comparison between muscle thickness and gastric motility would ideally require both measures to be evaluated under the same in vivo condition. However, there is no method currently established for this purpose.

The experimental data used in this paper only pertain to one type of diet with a fixed volume. The anatomical and motility features observed are representative of the postprandial state in healthy rats. However, the experimental and computational methods should be readily applicable to animals consuming different types of diets to assess their differential effects on gastric accommodation, motility, and emptying. The approach can also be applied to different groups of animals to evaluate the effects of various biological variables, pathophysiological vs. physiological conditions, or pharmacological vs. bioelectric treatment. As such, our approach provides a reliable means for preclinical studies of gastric motor functions.

There is no fundamental barrier to applying a similar approach to humans. Recent studies have demonstrated the feasibility of contrast-enhanced GI MRI in humans with natural ingredients [13], [14]. They used volumetric assessment or estimated gastric motility on the surface of the luminal content. By estimating the displacement of individual points on a surface surrounding the ingested food, [14] failed to directly consider the constraint that all the points, when moved, must bear a topology to span a smooth surface (i.e., manifoldness). Without satisfying this constraint, the motility is likely to be overestimated. Our approach can properly address this issue and merit its future applications to human GI MRI. Moreover, our approach provides a unique capability of performing group-level analysis and mapping gastric motility. This is particularly compelling for clinical applications. The surface motility patterns defined with normal subjects can serve as the baseline for comparison with disease populations. Their group-level differences can be mapped, thereby linking disease phenotypes to specific gastric locations or specific types of motor events.

